# Structure Basis for Inhibition of SARS-CoV-2 by the Feline Drug GC376

**DOI:** 10.1101/2020.06.07.138677

**Authors:** Xiaodong Luan, Weijuan Shang, Yifei Wang, Wanchao Yin, Yi Jiang, Siqin Feng, Yiyang Wang, Meixi Liu, Ruilin Zhou, Zhiyu Zhang, Feng Wang, Wang Cheng, Minqi Gao, Hui Wang, Wei Wu, Ran Tian, Zhuang Tian, Ye Jin, Hualiang Jiang, Leike Zhang, H. Eric Xu, Shuyang Zhang

**Affiliations:** School of Medicine, Tsinghua University, Haidian District, Beijing, China; Department of Cardiology, Peking Union Medical College Hospital, Peking Union Medical College and Chinese Academy of Medical Sciences, Beijing, China; Tsinghua-Peking Center for Life Sciences, Tsinghua University, Beijing, China; State Key Laboratory of Virology, Wuhan Institute of Virology, Center for Biosafety Mega-Science, Chinese Academy of Sciences, Wuhan, China; The CAS Key Laboratory of Receptor Research, Shanghai Institute of Materia Medica, Chinese Academy of Sciences, Shanghai 201203, China; Wuxi Biortus Biosciences Co. Ltd., 6 Dongsheng West Road, Jiangyin 214437, China; University of Chinese Academy of Sciences, Beijing 100049, China

## Abstract

The pandemic of SARS-CoV-2 coronavirus disease-2019 (COVID-19) caused by SARS-COV-2 continues to ravage many countries in the world. Mpro is an indispensable protein for viral translation in SARS-CoV-2 and a potential target in high-specificity anti-SARS-CoV-2 drug screening. In this study, to explore potential drugs for treating COVID-19, we elucidated the structure of SARS-CoV-2 Mpro and explored the interaction between Mpro and GC376, an antiviral drug used to treat a range of coronaviruses in Feline via inhibiting Mpro. The availability and safety of GC376 were proved by biochemical and cell experiments in vitro. We determined the structure of an important protein, Mpro, in SARS-CoV-2, and revealed the interaction of GC376 with the viral substrate and inhibition of the catalytic site of SARS-CoV-2 Mpro.

## Results and Discussions

The pandemic of SARS-CoV-2 coronavirus disease-2019 (COVID-19) caused by SARS-COV-2 continues to rampage across the world. The life cycle of SARS-CoV-2 requires the intact function of a 3C-like protease, also called the main protease (Mpro), encoded by the gene of nonstructural protein (NSP) 5. The function of Mpro is indispensable for viral replication and transcription and is therefore an attractive drug target against SARS-CoV-2. After viral infection in host cells, two polyproteins, pp1a (486kD) and pp1ab (790kD), are translated, and they are then cleaved by Mpro together with papain-like protease 2 (PL2^pro^). After the cleavage at 11 sites by Mpro and 3 sites by PL2^pro^, pp1a and pp1ab are processed to release a series of NSPs that have diverse functions to mediate viral replication and transcription^1^. Mpro also contains a distinct P2 Asn-specific substrate-binding pocket and has been reported to inhibit cellular translation by cleavage of poly(A)-binding protein^2, 3^.

To explore potential drugs for COVID-19 treatment, we aimed to solve the SARS-CoV-2 Mpro structure and study its interactions with potential drug molecules. Mpro was expressed in *Escherichia coli* cells and then purified via affinity and size-exclusion chromatography. The purified Mpro showed a monodispersed peak and a molecular weight of approximately 33.8 kD (Supplementary information, Fig S1). Mpro was crystallized and its structure was determined at 2.35 Å resolution. Like SARS-CoV, the SARS-CoV-2 Mpro forms a dimer with two protomers vertically packed with each other (Fig. 1a; Supplementary information, Table S1). The Mpro monomer contains three domains with domains I (residues 8-101) and II (residues 102-184) forming a barrel structure of six antiparallel β-strands (Fig. 1a, Supplementary information, Fig. S2a)^4^. Between domain I and II is a cleft that forms the catalytic dyad cys145-his41 and substrate binding site, which is comprised of the highly conserved substrate binding pockets (Supplementary information, Fig. S2b)^5, 6^. Domain III (residues 201-303) is comprised of a globular cluster of five antiparallel α-helices, which is involved in the dimerization of Mpro. Domain II and III are connected through a loop (residues 185-200) and domain I contains a stretch-out “N-finger” (NH2-terminal) that is inserted into domain II of the other protomer via interaction with F140 and G166 (Fig 1a, Supplementary information, Fig S2a)^6^. Meanwhile, the surface charge distribution in the active site of Mpro displays bi-polar distribution that is fit for the binding of peptide substrates (Fig. 1b).

**Figure 1.**
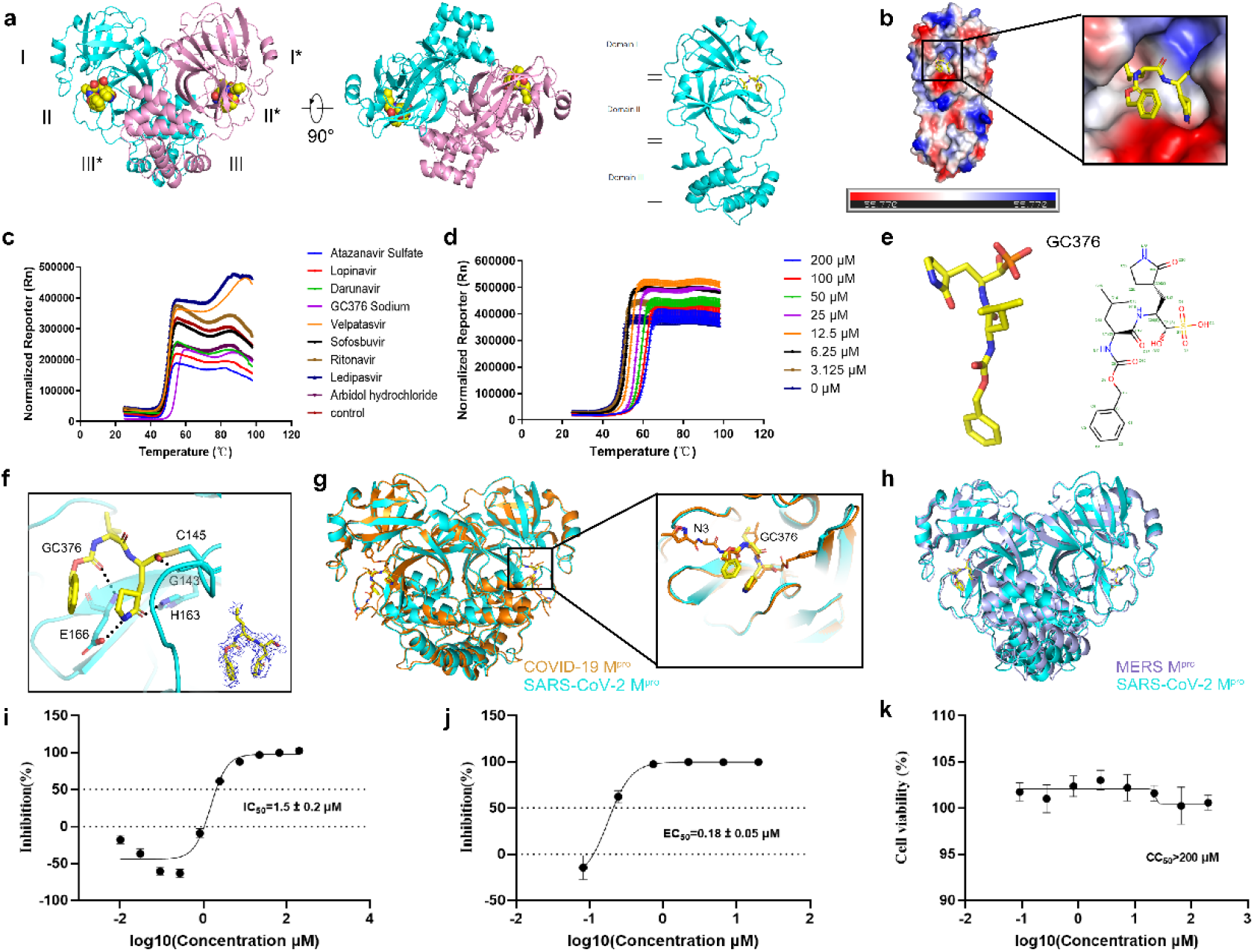
Structure of SARS-CoV-2 Mpro and activity of GC376 on Mpro. **a,** The overall structure of SARS-CoV-2 Mpro dimer in complex with GC376 in two different views and the structure of Mpro monomer. **b,** Electrostatic surface of SARS-CoV-2 Mpro. Blue: positive charge potential; Red: negative charge potential. The value ranges from −55.77 (red) to 0 (white) to 55.77 (blue). **c,** Thermal shift assay of SARS-CoV-2 Mpro with different drugs (Rn: normalized reporter signal). **d,** Thermal shift assay of SARS-CoV-2 Mpro with different concentrations of GC376. **e,** Chemical structure of GC376. **f,** Binding of GC376 (yellow sticks) to the SARS-CoV-2 Mpro pocket with the electron density map for GC376. **g,** Superposition of SARS-CoV-2 Mpro (cyan) and COVID-19 Mpro (orange) crystal structures. **h,** Comparison between GC376 bound SARS-CoV-2 Mpro (cyan) and MERS 3CLpro (mauve). **i,** IC50 of GC376 on the purified Mpro enzyme. **j,** EC50 of GC376 on inhibition of SARS-Cov-2 in Vero E6 cells. **k,** CC50 of GC376 on Vero E6 cells.

Several small molecules have been reported to effectively inhibit coronavirus Mpro^7^, and we performed thermal shift assay (TSA) to test the binding of each drug to SARS-CoV-2 Mpro. Each drug (20 μM) was incubated with Mpro and the melting profile was monitored based on the SYBRO orange reaction at 25-99 °C. GC376 shifted the melting curve of Mpro from 50.87°C to 55.23°C (+4.35°C), while other drugs show little effect (Fig. 1c, Supplementary information, Table S2), indicating the direct binding of GC376 to Mpro. To test if the process is titratable, TSA was repeated with different concentrations of GC376 (0-20 μM) and melting temperature of Mpro was increased with the increased doses of GC376 (Fig. 1d).

GC376 is a bisulfite adduct of a peptidyl derivative, containing leucine and proline with an additional phenylalanine ring (Fig. 1e). GC376 has been reported to be able to inhibit 3CL protease^8^, which gives GC376 a broad anti-viral spectrum, including human norovirus^9^. However, the drug now is currently used to treat animal coronavirus diseases, such as feline infectious peritonitis^10^. Because of its high binding affinity to SARS-CoV-2 Mpro verified by the TSA (+4.35°C), a pre-IND (Investigational New Drug application) about the usage of GC376 on COVID-19 treatment has been submitted to FDA^11^.

Structure of GC376-SARS-CoV-2 Mpro complex was solved through co-crystallization and GC376 was found to occupy the Mpro substrate binding site (Fig. 1f). The binding of GC376 to Mpro is stabilized by both hydrophilic and hydrophobic interactions. GC376 forms direct hydrogen bonds with G143, H163 and E166 of SARS-CoV-2 Mpro. The proline ring and leucine side chain of GC376 fit well into the Mpro S1 and S2 pockets, and form an extensive set of hydrophobic interactions with the conserved residues from the substrate binding pocket (Fig. 1f). In addition, after removing bisulfite group, GC376 can form covalent bond with Mpro Cys145, the catalytic residue, revealing its ability to directly block the catalytic dyad and its protease activity. This shows the potentiality of GC376 to act as a covalent inhibitor to prevent the binding and cleavage of the substrate. The structure of our GC376-SARS-CoV-2 Mpro complex was compared to the COVID-19 Mpro structure solved by Rao, et al^6^ (Fig. 1g) and MERS 3CLpro^12^ (Fig. 1h) respectively, and a good agreement can be observed. Comparison with COVID-19 Mpro shows high conservation of domain I and II, with small variable regions in domain III. Comparison with MERS Mpro shows the same localization of GC376, indicating that the inhibition of these viruses by GC376 is mediated through a similar mechanism.

To verify the effectiveness of GC376, we tested the ability of GC376 to inhibit SARS-CoV-2 Mpro enzymatic activity, which revealed a 50% inhibition concentration (IC50) of 1.5 μM (Fig. 1i). We also tested the ability of GC376 to inhibit SARS-CoV-2 infection in pre-seeded Vero E6 cells. GC376 showed a 50% maximal effect concentration (EC50) of 0.18 μM (Fig. 1j). In contrast, GC375 has a 50% cytotoxic concentration (CC50) of more than 200 μM, showing an excellent safety profile (Fig. 1k). Both structure analysis and *in vitro* tests by our studies indicate that GC376 is a relatively effective and safe drug candidate for SARS-CoV-2 by inhibiting Mpro. Further animal experiments and clinical trials are warranted for validating the potential use of GC376 for the treatment of COVID-19.

## Sources of support

This work was supported by CAMS Innovation Fund for Medical Sciences (CIFMS) (No. 2016-I2M-3-011, 2016-I2M-1-002), CAMS Innovation Fund for Medical Sciences No. (2020-I2M-CoV19-001). “13th Five-Year” National Science and Technology Major Project for New Drugs (No: 2019ZX09734001-002), and Tsinghua University-Peking University Center for Life Sciences (No. 045-160321001) to S.Y.Z.; and the National Key R&D Programs of China 2018YFA0507002, Shanghai Municipal Science and Technology Major Project 2019SHZDZX02 and XDB08020303 to H.E.X. We appreciate the help from Sangon Biotech Co. Ltd for DNA synthesis, and Vazyme Biotech Co. Ltd for offering ClonExpress II One Step Cloning Kit.

## Author Contribution

X.L. performed the protein purification and structure analysis of Mpro and wrote the paper under the supervision of S. Z.; W.S. performed cell experiments under the supervision of L. Z.; F. W.; W.C., M.G. assisted in the work of X. L.; Y.W., M. L., and R.Z. participated in the manuscript writing and figure preparation; S. Z., H.J. and H.E.X. helped design the study and wrote the paper.

## Supplementary Information

### Materials and methods

#### Protein Expression and Purification

The gene encoding SARS-CoV2 Mpro containing a histidine-tag followed by the TEV protease cleavage site was cloned into pET28a plasmid vector. The construct was transformed into *Escherichia coli* BL21 (DE3) cells and cultured in LB medium. 0.5 mM IPTG was added to induce expression at 15 °C for 16 hours. Then the cells were broken up by High Pressure Homogenizer and centrifuged at 18000 rpm for 60 min. The supernatant was purified by affinity chromatography (His FF) and the His tag of Mpro was eluted by cleavage buffer (50mM Tris-HCl (pH 8.0), 500mM NaCl, 5% glycerol, 300mM imidazole). Then the protein underwant size-exclusion chromatography (Superdex 200 Increase 10/300 GL). TEV protease was added and incubated over night at 4 °C. The Mpro was further purified by ion affinity chromatography (His HP) and Size-exclusion chromatography (HiLoad 16/600 Superdex 200 pg). Finally, quality control of Mpro was conducted through SDS-PAGE, analytical size-exclusion chromatograph (Superdex 200 Increase 5/150 GL) and liquid chromatography-mass spectrometry (LC-MS).

#### Crystallization of Mpro

SARS-CoV2 Mpro was incubated on ice for 2h after the 1mM GC376 Sodium added. 500 nL protein (10.81mg/ml) and 500 nL reservoir were added. The complex was crystallized by the hanging drop vapor diffusion method at 20°C. The best crystals were grown with 0.2M Lithium chloride, 0.1M Hepes pH 7, 20% w/v PEG 6000 buffer, with a protein concentration of 5 mg/ml. Crystals were obtained after 3 days of growth.

#### Data collection and structure determination

X-ray diffraction data were collected at (CLS in Canada) at beam line (CLSI BEAMLINE 08ID-1) using a wavelength of (0.97949) Å. The Mpro structure was determined by molecular replacement (MR) with (phaser), using the X-Ray Crystal Structure of the SARS Coronavirus Main Protease as a template (PDB code: 1q2w). The model was then manually adjusted by COOT.

#### Thermal Shift Assay

Thermal shift assay was performed in Applied Biosystems 7500 fast real time system. For each reaction, 5 μL of compound working solution was added to 96-well RT-PCR plate (AXYGEN, USA). Then 10 μL Mpro purified protein (0.5 mg/ml) working solution was added to each well. The different compounds were incubated with protein for 1 hour at 4°C. After incubation, 5 μL dye SYPRO Orange working solution was added to each well. The reaction was run in an ABI 7500 fast real-time PCR system with a ramp rate of 1°C per minute from 25°C and ending at 99°C.

#### Viral Infection Experiments

Vero E6 cell line was obtained from American Type Culture Collection (ATCC) and maintained in minimum Eagle’s medium (MEM, Gibco Invitrogen) supplemented with 10% fetal bovine serum (FBS, Gibco Invitrogen), 1% antibiotic/antimycotic (Gibco Invitrogen), at 37°C in a humidified 5% CO_2_ incubator. SARS-CoV-2 (nCoV-2019BetaCoV/Wuhan/WIV04/2019) was propagated in Vero E6 cells^1^, and viral titer was determined by 50% tissue culture infective dose (TCID50) as described in previous study^2^. All the infection experiments were performed in a biosafety level-3 (BSL-3) laboratory.

#### The IC50 assay of GC376 sodium for 3CLpro

Three-fold serial dilute the GC376 sodium stock solution (10mM, DMSO) with DMSO to get the 50x compound solution, whose concentrations range from 0.5-10000μM (10 doses). 8.3 fold dilute 50x compound solution with pure water to prepare 6x GC376 sodium solution containing 12% DMSO, whose concentrations range from 0.01-200μM. Then use 1.2x reaction buffer (24mM HEPES pH7.0, 19.2mM NaCl, 2.4mM DTT, 0.012% Triton-X100) to dilute 3CLpro to 375nM and peptide Dabcyl-KTSAVLQSGFRKME-Edans (10mM, DMSO) to 36μM, to get 1.5x 3CLpro solution and 6x substrate solution respectively. Add 10μL 6x substrate (36μM), 40μL 1.5x 3CLpro (375nM) and 10μL 6x compound solution (different concentration) to each well in 384-well Micro plate (Corning, #3575). Make sure final DMSO concentration is 2% in each well. After adding reagents, centrifuge plate and vortex. Immediately after vortex, read the signal for 30min via TECAN M1000 plate reader through kinetic model. The instrument settings are shown in table S3. Data was analyzed via GraphPad Prism 6.0.

#### Evaluation of antiviral activities of the test compounds

Vero E6 pre-seeded in 48-well dish (1 x 10^5^ cells/well) were treated with the different concentration of the indicated compounds for 1 hours and infected with SARS-CoV-2 at a MOI of 0.05. Two hours later, the virus-drug mixture was removed and cells were cultured with drug containing medium. At 24 hours p.i, the cell supernatant was collected and lysed. The viral RNA extraction and quantitative real time PCR (RT-PCR) analysis was described in previous study^2^.

#### Evaluation of the cytotoxicity of the test compounds

Vero E6 pre-seeded in 96-well dish (5 x 10^5^ cells/well) were treated with different concentration of the indicated compounds, and 24 hours later, the relative numbers of surviving cells were measured with cell counting kit-8 (GK10001, GLPBIO) according to the manufacturer’s instructions.

---

**Table S1.** Data collection and structure refinement statistics.

| **Data collection** |  |
| --- | --- |
| Space group | C 2 |
| Resolution (Å) | 50.00-2.35 (2.43-2.35) |
| Cell parameters |  |
| a, b, c (Å) | 114.303, 53.874, 45.233 |
| $\alpha$, $\beta$, $\gamma$ (°) | 90.000, 101.576, 90.000 |
| Completeness (%) | 98.9(99.4) |
| Total/Unique reflections | 33738(3406)/ 11227(1112) |
| Redundancy | 3.0(3.1) |
| Rmerge (%) | 5.5(36.8) |
| Rmeans# | 6.6(44.5) |
| Mean I/$\sigma$ | 11.5(2.6) |
| CC 1/2 | 0.998(0.883) |
| **Refinement** |  |
| Resolution range (Å) | 44.31 -2.35 (2.43-2.35) |
| No. reflections | 10663(743) |
| Wilson B factor, Å<sup>2</sup> | 29.957 |
| Rcrystal % | 0.2143 |
| Rfree % | 0.2707 |
| No. of non-H atoms |  |
| Protein | 2331 |
| Ligand/ion | 29 |
| Water | 35 |
| B factors (Å<sup>2</sup>) |  |
| Protein | 44.0 |
| Ligand/ion | 63.3 |
| Water | 36.8 |
| R.m.s. deviations |  |
| Bonds (Å) | 0.004 |
| Angles (°) | 1.401 |
| Ramachandran plot |  |
| Most favoured (%) | 96 |
| Allowed (%) | 4 |
| Outlier(%) | 0 |
| Clash score | 5 |

**Table S2.** Different compounds added for thermal shift assay and the results of melting temperature.

| NO. | Concentration | Compound | Tm1 | Tm2 | Tm3 | Tm Mean (°C) | Δ Tm (°C) |
| --- | --- | --- | --- | --- | --- | --- | --- |
| 1 | 20 uM | Atazanavir Sulfate | 50.56 | 50.56 | 50.37 | 50.50 | -0.38 |
| 2 |  | Lopinavir | 50.94 | 50.75 | 50.75 | 50.81 | -0.06 |
| 3 |  | Darunavir | 51.13 | 51.13 | 51.13 | 51.13 | 0.25 |
| 4 |  | GC376 Sodium | 55.29 | 55.29 | 55.1 | 55.23 | 4.35 |
| 5 |  | Velpatasvir | 50.94 | 50.94 | 50.94 | 50.94 | 0.06 |
| 6 |  | Sofosbuvir | 51.13 | 51.13 | 50.94 | 51.07 | 0.19 |
| 7 |  | Ritonavir | 50.56 | 50.56 | 50.37 | 50.50 | -0.38 |
| 8 |  | Ledipasvir | 50.37 | 50.37 | 50.18 | 50.31 | -0.57 |
| 9 |  | Arbidol hydrochloride | 50.18 | 50.56 | 50.75 | 50.50 | -0.38 |
| 10 |  | control | 50.94 | 50.94 | 50.75 | 50.88 | 0.00 |

**Table S3.** Instrument Setting of TECAN M1000 plate reader.

| Mode | Fluorescence Top Reading |
| --- | --- |
| Excitation Wavelength | 340nm |
| Emission Wavelength | 535nm |
| Excitation Bandwidth | 5nm |
| Emission Bandwidth | 20nm |
| Gain | 110 Manual |
| Number of Flashes | 50 |
| Flash Frequency | 400Hz |
| Integration Time | 20μs |
| Lag Time | 0μs |
| Settle Time | 0ms |
| Z-Position (Manual) | 23792μm |

**Figure. S1.**
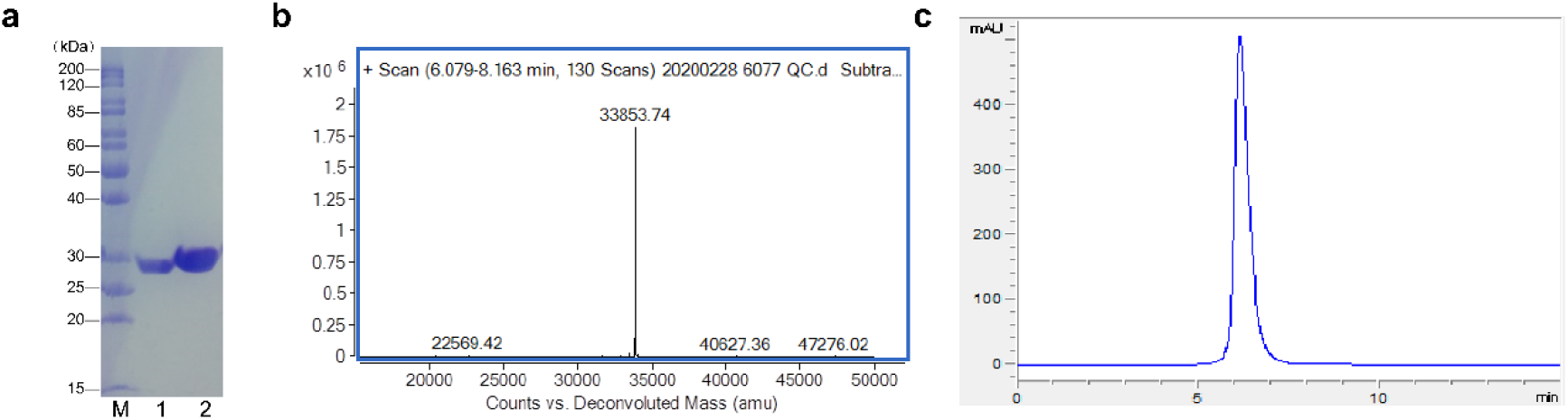
Protein expression and purification of SARS-CoV-2 Mpro. **a,** SDS-PAGE result of SARS-CoV-2 Mpro. **b,** Analytical size-exclusion chromatography (SEC) of SARS-CoV-2 Mpro. **c,** Liquid chromatography-Mass Spectrometry (LC-MS) of SARS-CoV-2 Mpro.

**Figure. S2.**
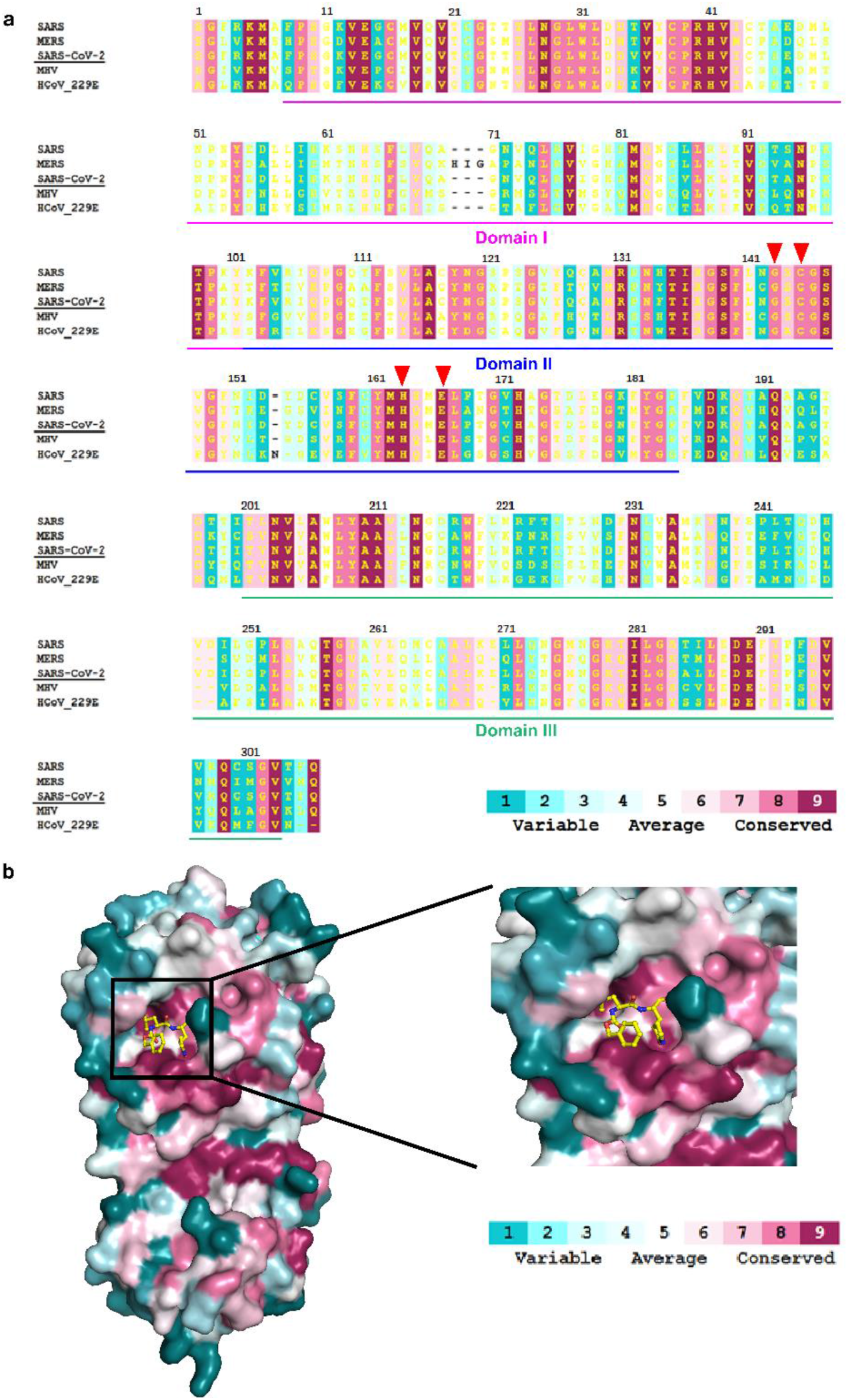
The conservatism of SARS-CoV-2 Mpro. **a,** The amino acid sequence of SARS-CoV-2, SARS-CoV, MERS-CoV, MHV and HCoV-229E were compared and the conservatism was shown by different color. The sequence of domain I, II and III were marked. The triangles indicate the amino acids interact with GC376. **b,** The SARS-CoV-2 Mpro structure and the sites interact with GC376 were shown, and the color represents the conservatism.

## Notes

### Competing Interest Statement

The authors have declared no competing interest.

